# Genomic insights into the karyotypic radiation of a narrow endemic holocentric plant *Carex helodes*

**DOI:** 10.64898/2026.07.14.738159

**Authors:** Inés Gómez-Ramos, Rogelio Sánchez-Villegas, Ashwini V. Mohan, Camille Cornet, André Marques, Enrique Maguilla, Santiago Martín-Bravo, Kay Lucek, Marcial Escudero

## Abstract

Holocentric chromosomes allow rapid genome changes through chromosomal rearrangements such as fissions, fusions, inversions or translocations. The plant genus *Carex* shows one of the highest rates of karyotypic evolution among holocentric organisms. We studied the genomic patterns underlying chromosomal rearrangements in the karyotypic radiation of the narrow endemic species *Carex helodes* (2n = 68-75).

Comparing genome assemblies of *C. helodes* from the two karyologically distinct extremes of its European distribution, revealed a striking number of eight chromosomal rearrangements including fusions, translocations and inversions. Genomic breakpoints are gene-poor and TE-rich, corroborating findings in other species and suggesting common genomic characteristics that facilitate the evolution and establishment of chromosomal rearrangements. We identified a chromosomal inversion exhibiting patterns of purifying selection and enrichment in functional genes that potentially mediate rearrangement tolerance. Conversely, another inversion displayed elevated sequence divergence and enrichment in response to temperature stress and phosphate limitation, matching key environmental variables that differ between the study localities.

The establishment of chromosomal rearrangements along *Carex helodes* European populations was likely driven by demographic bottlenecks and distinct genomic features at breakpoints. Our findings provide preliminary evidence on the rearrangement role in population differentiation either as reproductive barriers or as genomic islands of differentiation.

## Introduction

The role of chromosomal rearrangements (CRs, encompassing fusions, fissions, inversions, duplications, insertions, deletions and translocations) in eukaryote evolution is a long standing debate [1–3]. While some authors highlight their importance [1,3,4], others consider them to be largely incidental [5–7]. In plants, dysploid CRs are more frequent and significant for evolution than expected compared to polyploidy [8–10], despite their potential deleteriousness e.g. altering gene function or expression [11–13] and causing difficulties for chromosome pairing during meiosis in heterokaryotypes [1,14]. Despite this potential deleteriousness, CRs offer an opportunity for speciation when they cause hybrid dysfunction, and therefore, act as a barrier to gene flow [1,3]. CRs may also contribute to speciation through recombination suppression [1,2,15], whereby recombination between chromosomes carrying different CRs is reduced, promoting divergence and decreasing introgression in those regions. If CRs encompass multiple co-adapted genes that are physically coupled, they can be considered as a single supergene [16]. Suppressed-recombination speciation is especially efficient when the rearranged regions contain genes that are key for both reproductive isolation and ecological adaptation [17].

The ability for underdominant CRs to establish within populations and/or propagate to new areas is influenced by population dynamics [1,18]. For example, within small populations where drift is strong compared to selection, a new CR is more likely to establish even under strong underdominance [19,20]. Also, partial reproductive barriers within a population may allow for some genetic structure and for underdominant CRs to establish within subpopulations. Inbreeding, vegetative propagation and long life cycles may similarly help the establishment of new CRs within a population [21,22]. Some of these features can also promote the colonization of new areas, where drift is higher and, therefore, the establishment of new CRs could be easier. Heterogeneous environments can facilitate the persistence of new CRs within a different ecological niche, as locally adapted alleles may arise and become fixed within rearranged areas where recombination is suppressed [23,24].

Similar CRs often arise repeatedly over the evolutionary history of lineages [11,25–27]. This recurrent aspect of CRs is reflected by the increased degree of conserved syntenic blocks in taxonomic groups with high CR rates, where CRs are constrained to regions with certain genomic features [25]. Such genomic hotspots for CRs have been reported for both plants [25,28], Lepidoptera [26] and mammals [29,30]. For example, CR hotspots are enriched for repetitive regions that are more prone to non-allelic homologous recombination (NAHR) [31,32]. Breakpoints also tend to be outside of gene rich regions, likely because CRs could break gene sequence or separate it from regulatory regions [25,33].

While the mechanisms underlying CR formation are conserved throughout eukaryotes [34,35], their evolutionary impact largely varies between different clades. The importance of underdominance for CRs has been suggested to be lower for species with holocentric chromosomes, where kinetochore activity is distributed throughout each chromosome instead of being located in one region [36–38]. Contrary to monocentric species, fissions and fusions are expected to be more tolerated in holocentric organisms [39,40] leading to increased rates of CR evolution with frequent meiotic irregularities [39,41,42]. Some holocentric clades have distinct genomic features such as a more uniform GC content and gene distribution along their chromosomes [43–46], which is otherwise associated with the distance to the centromere in monocentric organisms [47–49]. In summary, the principles of CR evolution may differ between holocentric and monocentric clades [50].

One of the best studied holocentric lineages is the genus *Carex* L. (Cyperaceae; commonly known as true sedges), exhibiting high rates of CRs [37,41,51–55], which has been suggested to underlie its striking species richness (ca. 2000 sp.) [37,41,52,56]. This has been confirmed by recent phylogenetic macroevolutionary studies [57,58]. Significant differences in karyotype evolution occur both across and within different *Carex* clades [57]. For example, within Spirostachyae [54], some species have a stable chromosome number and others showing large ranges of intraspecific chromosome number variation (*Carex laevigata* Smith 2n = 69 - 84 or *C. helodes* Link 2n = 68 - 75). In fact, *C. helodes* even shows chromosome number variation between individuals with no genetic differentiation at the Internal Transcribed Spacer (ITS) DNA region or microsatellites, suggesting a recent origin [59,60].

Here, we investigate the genomic basis of the remarkable karyotypic diversity observed across the European range of the Ibero-African narrow endemic sedge *Carex helodes* [60]. Firstly, we characterize CRs underlying changes in chromosome number and estimate the number of CRs through synteny analyses between two individuals sampled from two populations from opposite extremes of the species’ European range. We then test the hypothesis that CRs are enriched for genomic features that either facilitate their emergence and/or limit their deleteriousness, by comparing the genomic characteristics of breakpoints and rearranged chromosomes with those of conserved regions. We further evaluate the impact of CRs on gene evolution in inverted regions and areas flanking breakpoints, where recombination is expected to be suppressed [1,15]. Finally, the functions of these genes are assessed through gene ontology analysis to infer the potential impact of CRs.

## Methods

### (a) Study species

*Carex helodes* Link, a member of the *Carex* subgenus *Carex* in the Cyperaceae family [61], is a diploid, wind-pollinated, perennial herb with caespitose habit (without creeping rhizomes), rough upper leaf surface and high numbers of male (1–4[–7]) and androgynous spikes. It is endemic to southern Portugal, southwestern Spain and northern Morocco [62].

The *C. helodes* karyotype varies from 2n = 74 in Morocco (exceptionally 2n = 75 in Portugal) to 2n=68 in Aznalcollar, Spain [54,60,62]. An intrapopulation cytological study demonstrated that karyotypical diversity was even high within populations, with four different chromosome numbers (2n=68, 69, 70, 72) and six different karyotypes [60] suggesting the presence of multiple CRs, with 2n=70=35^II^ being by far the most common karyotype. Despite this great karyotypic diversity, all individuals in the population were genetically identical based on 34 microsatellites that were variable for the species outside this population [60]. Former genetic studies on *C. helodes* [60,62] point to Portuguese populations as being ancestral and genetically diverse while Spanish populations evolved more recently and have a much lower genetic diversity, consistent with a west-to-east expansion from southern Portugal to southwestern Spain involving multiple genetic bottlenecks and drift [60].

### (b) Plant material, library construction and sequencing

During spring 2019, we collected two individuals from two populations of *C. helodes,* located at the west-eastern limits of the species distribution area within Europe. One individual was sampled in Aznalcollar (Seville, Spain; 37° 34’ N, 6° 21’ W) and the other in Caldas de Monchique (Algarve, Portugal; 37° 16’ N, 8° 31’ W), from now on referred to as AZN and MON, respectively. The populations represent the two extremes of the karyotype gradient for this species in Europe (figure 1*a*). The living plants were kept at the greenhouse of the University Pablo de Olavide, Spain. Experimental crosses and chromosome counts of parentals and F1 were performed (see electronic supplementary materials, Supplementary Methods, section (a)).

**Figure 1:**
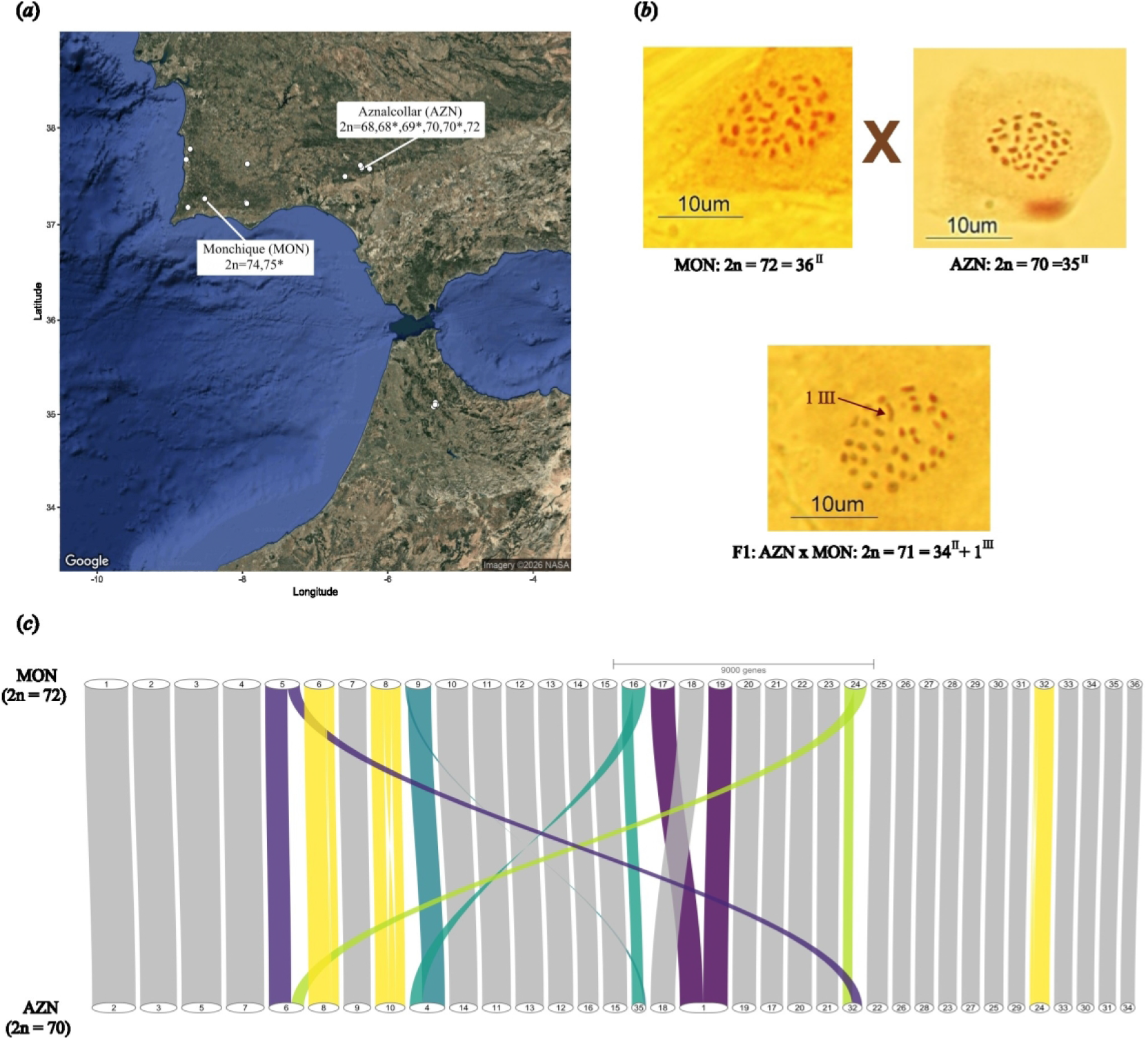
Geographic locations, meiotic configurations, and chromosomal synteny between *Carex helodes* parental genomes. (*a*) Satellite image of southern Spain and Portugal and northern Morocco with *Carex helodes* locations indicated [60,62,128]. Those included in our study are indicated, including previously reported chromosome numbers [60,62]. (*b*) Meiotic configurations of *C. helodes* parentals, MON (Caldas de Monchique, Algarve, Portugal) and AZN (Aznalcollar, Spain), and their F1 hybrid. Aberrant configurations are indicated by arrows. (*c*) Synteny plot generated with GeneSpace showing conserved gene collinearity and rearrangements between the genome assemblies of the parentals MON and AZN. Chromosomes are scaled by gene rank order Chromosomes with inversions are highlighted in yellow and interchromosomal rearrangements (translocations, fusions and fissions) in purple and green tones.

Fresh leaves were harvested and flash frozen with liquid nitrogen before storage at -80°C. High-molecular weight DNA extraction and long-read HiFi sequencing on a PacBio Revio was outsourced to Novogene (Cambridge, UK). Hi-C was generated from fresh young leaf tissue. Firstly, nuclei extraction was carried out using the CelLytic™ PN Isolation/Extraction Kit from Merck. Then, DNA crosslinking, ligation, amplification, and library preparation was done using the Arima Hi-C high-throughput kit and a modified protocol for plant tissue. Finally, the prepared Hi-C libraries were paired-end sequenced on an Illumina Novaseq X (Illumina Inc., San Diego, CA, USA).

### (c) Genome assembly

HiFi data quality was assessed through NanoPlot v1.20.0 [63] and a k-mer histogram performed with kmer-jellyfish 2.3.1 [64] and GenomeScope 1.0 [65]. The length, coverage and quality of the long reads were considered adequate for both genomes.

Hi-Fi data was assembled using HifiASM 0.25.0 [66] obtaining contig level primary and alternate assemblies. Hi-C data was used in this step to correct sequencing and assembly errors. Duplicates were purged through a preliminary HifiASM purge and a further analysis with purge_dups 1.2.6 [67]. Scaffolding was performed using Hi-C data by mapping the Hi-C reads against each assembly following the Arima Hi-C mapping pipeline [68]. Then, scaffolding was performed with YaHS 1.2a.2 [69] with later manual curation performed in JuiceBox 2.15 [70] to correct misalignments between scaffolds. At each step of the assembly, quality controls were performed addressing i) assembly contiguity with gfastats 1.3.11 [71], ii) assembly completeness based on BUSCO genes from the embryophyta_odb10 and poales_odb10 databases using CompleASM 0.2.27 [72] and iii) with haplotype specific analysis based on kmer using Merqury 1.3 [73]. Contaminant scaffolds (e.g. organellar sequences, microbial sequences, or short low-coverage fragments) were identified using Tiara 1.0.3 [74] and removed. Finally, telomeric regions were identified and visualized to evaluate how many scaffolds were telomere-to-telomere using tidk v0.2.7 [75].

### (d) Gene annotation, synteny analysis and CRs identification

Gene annotation was performed with Braker3 v3.0.8 [79], which combines two methods: an homologous search using protein data from poales_odb10 [78] via the –Busco_lineage option and an *ab initio* approach. Annotation completeness has been assessed with BUSCO genes from the embryophyta_odb10 and poales_odb10 databases using CompleASM [72] using the option *analyze*. Prior to gene annotation, repetitive elements were softmasked with EarlGrey v4.4.5 [79] (further details are shown in electronic supplementary materials, Supplementary methods, section (b)).

Synteny analysis between the two genomes was performed with GENESPACE v.1.2.3 [81], using default parameters. Rearrangement breakpoints were defined as genomic regions which were not included in syntenic blocks. Rearrangements were manually explored with the Interactive Genomic Viewer [81] to ensure that there were no false rearrangements, e.g. due to assembly errors of repetitive regions close to telomeres [82]. CR validation was performed through two methods: reciprocal Hi-C cross-mapping and long read split end support (see supplementary materials, Supplementary methods). The possibility of heterozygosity for the detected inversions was also assessed (see supplementary materials, Supplementary methods, section (c)).

### (e) Characterization of conserved versus rearranged genomic regions

Conserved (collinear) regions and breakpoints were compared for gene density, GC content, total repeat content and repetitive elements families. The most common tandem repeat was separately analysed, as it is a possible candidate for holocentromeres (Table S1 in [25]). In addition, comparisons for the same genomic features were made between collinear chromosomes and rearranged ones. Prior to the comparative analyses, the relationships between genomic variables were assessed through pairwise correlation with the R package ggcorrplot v1.4.1 [83] using Spearman’s rank correlation coefficient.

Linear and generalized linear models were first performed with each genomic characteristic. Each genomic feature was defined within 100 kb windows throughout the whole genome for AZN and MON with the ‘*tileGenome*’ function from the GenomicRanges R package 1.34.1 [84]. The basic model formula was: Genomic variable ∼ (RC + BP:RC)*DC, which was run for each genomic feature separately. The independent variables include: RC (does the window belong to a rearranged chromosome?), BP (does the window include a rearrangement breakpoint?) and DC (window’s distance to the center of the chromosome). The variable BP is nested within RC. Chromosome identity was included as a random factor for a genomic characteristic model when it increased AIC and reduced residuals variance. Different R packages were used to perform these models: *‘lm’* and ‘*glm*’ function from stats for linear models, ‘*boxcox’* transformation from the package MASS 7.3-65 [85], ‘*glm.nb’* from the MASS package for negative binomial GLMs, ‘*lmer’* function from lme4 1.1-37 [86] for linear models with random effects and ‘*glmmTMB*’ from glmmTMB 1.1.9 for GLM with random effects [87]. The final model and the transformation used for each genomic characteristic are shown in table S4 (see electronic supplementary material). Chromosome size was compared between rearranged and collinear chromosomes, using Wilcoxon rank-sum and Student’s t-tests in R, ensuring that RC effects were not confounded by differences in chromosome size.

Breakpoint characterization was performed with ‘*permTest’* from the regioneR package 1.30 [88], which implements an enrichment analysis based on permutation tests through randomizing genome areas. The robustness does not depend on the number of breakpoints and is able to detect effects at finer scales. The number of overlaps (‘numOverlaps’) was used as the evaluation statistic, except for GC content whose evaluation statistic was defined as ‘meanInRegions’. Randomization was performed with ‘*randomizeRegions*’ from the same package with 1000 permutations. Breakpoints were compared against regions within the same chromosome and telomeres (defined as regions within 10kb of the chromosome ends) were excluded by masking them. Another permutation was performed dividing chromosomes into subtelomeric (10% most terminal sections of a chromosome) and interstitial sections.

Breakpoints were compared against regions of the same chromosome within the same genomic section using the option ‘universe’ in regioneR. Section limits were assessed based on the average point where GC content stabilizes from the telomeres. The magnitude of the enrichment was quantified by dividing the observed value by the median of the permuted distribution (*fold enrichment*). For zero-inflated variables (>20% of zeros) pseudocounts were used for the fold enrichment calculation: (observed value +1) / (mean permuted value + 1).

### (f) Divergence and selection in rearrangements

To assess patterns of molecular evolution associated with rearranged regions, sequence divergence between the two genomes was compared between rearranged and collinear regions of rearranged chromosomes. In this analysis the MON genome was used as reference, as is likely to be the ancestral population given its demographic history [60]. Because recombination can be suppressed within inversions and regions flanking other CRs [15,89,90], divergence—whether driven by genetic drift or selection—is expected to be higher within these regions. In this analysis rearranged regions included: (i) regions inside inversions, and (ii) regions flanking translocations and fission/fusion breakpoints (200 kb to each side of the non-collinear region). For the fusions, the rearranged regions were defined as 200 kb from the two chromosome ends (chr17 and chr19) and it was compared to the rest of the two chromosomes. DNA and amino acid sequences of genes located in rearranged chromosomes of the MON individual and their orthologous counterparts in AZN (obtained from Genespace) were aligned using MAFFT v7 [91]. Each coding sequence (CDS) was analyzed individually to maximize alignment accuracy. Alignments with <120 bp and/or >10% gaps were filtered out. Codon alignment distinguishing between synonymous and non-synonymous mutations was performed using pal2nal 14.1 [92]. Next, synonymous (dS) and non-synonymous (dN) substitution rates and their ratio (ω=dN/dS) were estimated through the ‘*codeml*’ function implemented in PAML [93], under the evolutionary Model 0, which assumes a constant ω throughout the tree. CDS with high (>2) dN or dS values were discarded, because when multiple substitutions occur in the same CDS, saturation occurs, compromising the rate’s reliability. Finally, substitution rates and their ratios were compared between rearranged regions and the rest of the rearranged chromosome. The ratio ω is a measure of selective pressure indicating if there is neutral evolution (ω ≈ 1), positive (ω > 1) or purifying selection (ω < 1). Ratio ω comparisons tested if selective pressure was affected by the rearrangements. The statistical significance of the differences was evaluated through enrichment analyses performed with regioneR [88]. Regions of the same size as the rearranged areas were sampled across the rest of the rearranged chromosome after telomeres were masked (10 kb from each extreme of the chromosome). Aggregated dN, dS, and ω values were calculated over 1,000 iterations.

### (g) Gene ontology within rearranged regions

Gene ontology (GO) enrichment analyses within rearranged regions (as defined before) were performed using topGO 2.50.0 [94] in R based on the annotated gene set obtained from Braker3 [79]. A custom gene-to-GO mapping was generated using eggNOG-mapper v2.1.12 [95,96]. Enrichment for the three primary Gene Ontology domains—Biological Process (BP), Cellular Component (CC), and Molecular Function (MF)—was assessed using classic Fisher’s exact tests implemented in topGO. GO-annotated genes within rearranged regions were considered the foreground and the rest of GO annotated genes in the genome as the background. GO-enrichment test were performed for all CRs combined as well as separately per CR for the MON genome Redundant BP GO terms were clustered using GOSemSim 2.24.0 [97] based on the *Arabidopsis thaliana* annotation database (<u>org.At.tair.db</u>) for visualization.

## Results

### (a) Genome assembly and annotation

*Carex helodes* long read sequencing yielded 60 Gb of data from 8.33 million HiFi reads representing a coverage of 42.4x for Aznalcollar (AZN) and 72 Gb of data from 14.1 million reads for a coverage of 49.2x for Monchique (MON). Hi-C data accounted for 145 Gb from 392 million reads for AZN and 204 Gb from 569 million reads for MON.

Our final assemblies were 346.74 Mb in 35 chromosome-scale pseudomolecules with more than 99.98% of the bases in the assembly for AZN and 348.51 Mb in 36 chromosome-scale pseudomolecules with more than 99.77% of the bases in the assembly for MON. The assemblies had a scaffold N50 of 10.12 Mb and 9.69 Mb for AZN and MON, respectively. More than half of the chromosome-scale pseudomolecules had telomeric repeats at both chromosome ends (see electronic supplementary materials, figure S1). BUSCO analysis identified around 93 % of complete single-copy genes and 4 % of missing genes for both genomes when using the lineage Embryophyta as a reference (see electronic supplementary materials, table S1). However, when using the more complete gene set of the Poales lineage, the proportion of missing genes rose to 17 % for AZN and 18 % for MON, which is consistent with other high quality *Carex* genomes [98,99] as the reference gene set contains proteins unique to Poaceae.Parental genomes were highly homozygous (0.23% and 0.26% heterozygosity for AZN and MON, respectively, according to GenomeScope [65]), consistent with other *Carex* genomes where clonal reproduction is not prevalent and repeat content is relatively high with 50.20% of repetitive DNA in AZN and 50.38% for MON.

Gene annotation yielded 30,745 genes for AZN and 31,224 for MON. Missing genes did not increase after gene annotation based on BUSCO analysis with Poales as the reference dataset and only by 2 % when Embryophyta is the reference used (see electronic supplementary material, table S1).

### (b) Synteny analysis: A surprising number of rearrangements

The synteny analysis performed with Genespace showed three inversions (Inv_chr6, Inv_chr8 and Inv-chr32; rearrangements were named according to the more ancestral MON genome [60]), 4 translocations (Trans_chr5, Trans_chr9, Trans_chr16 and Trans_chr24) and one fusion (Fusion-chr17 and fusion-chr19), which results in the largest AZN chromosome (Figure 1*c*). These CRs were confirmed through crossed Hi-C contact maps (see electronic supplementary materials, figure S2). Long–read split-read support analyses were able to adequately predict breakends for most CR breakpoints in at least one of the genomes (see electronic supplementary materials, table S2). The inversion in chr32 was heterozygous in the MON genome in accordance to both Hi-C contact maps (see electronic supplementary materials, figure S3) and long-read split-read support (76 out of 130 support the inversion, see electronic supplementary material, table S2).

### (c) Germination success and cytogenetics

F_1_ and F_2_ germination successes were 86% (obtained from one cross) and 60% (mean: 60%, sd: 0.28, number of crosses: 9), respectively. The cytogenetic experiments, performed on the same parentals as the synteny analysis, revealed chromosome numbers of 2n = 36^II^ = 72 for the MON parent and 2n = 35^II^ = 70 for the AZN parent, both displayed exclusively bivalent formations. In the five meioses analyzed for these two F₁ individuals, all showed the same configuration. Specifically, the F₁ individuals showed intermediate chromosome numbers (2n = 34^II^+1^III^ = 71) with 34 bivalents and one trivalent (figure 1*b*).

## (d) Genomic differences between rearranged and conserved areas

Both genomes were divided in 100kb windows resulting in 3467 windows for AZN and 3486 windows for MON, with 18 windows containing rearrangement breakpoints for AZN and 15 for MON. Most genomic features varied among chromosomes, with the exception for tandem repeat clusters (TRCs) and the most common TRC (electronic supplementary material, table S4, figure S4). The best predictor for all genomic features, except gene, LINE and SINE content, was the position of the window within a chromosome (DC) (electronic supplementary material, table S4, figure S4). Overall TE proportion, TEs subtypes (except LINEs and SINEs) and GC content increased towards the telomeres, while unknown repeats followed the opposite pattern, as demonstrated by the negative estimate of the DC effect for unknown repeats. This negative correlation between unknown repeats and GC content is reflected in the autocorrelation analysis (see electronic supplementary materials, figure S6), the highest correlation between variables that do not derive from each other.

The genomic composition differed significantly between rearranged and conserved chromosomes (figure 2b, electronic supplementary material, table S4, figure S4): rearranged chromosomes displayed less LINEs for both genomes and lower GC content for the MON genome. These differences were not explained by chromosome size (AZN: P (t test)=0. 29; AZN: P (Wilcoxon)=0.36; MON: P (t test)=0.47; MON: P (Wilcoxon)=0.27). If the interaction between RC and DC is considered, rearranged chromosomes have a lower TE density towards the telomeres. LTRs and within those Ty3/Gypsy had also significant effects in the same direction. However, this was only significant for AZN, while for MON only DNA transposons were scarcer in rearranged chromosomes near the telomeres.

**Figure 2:**
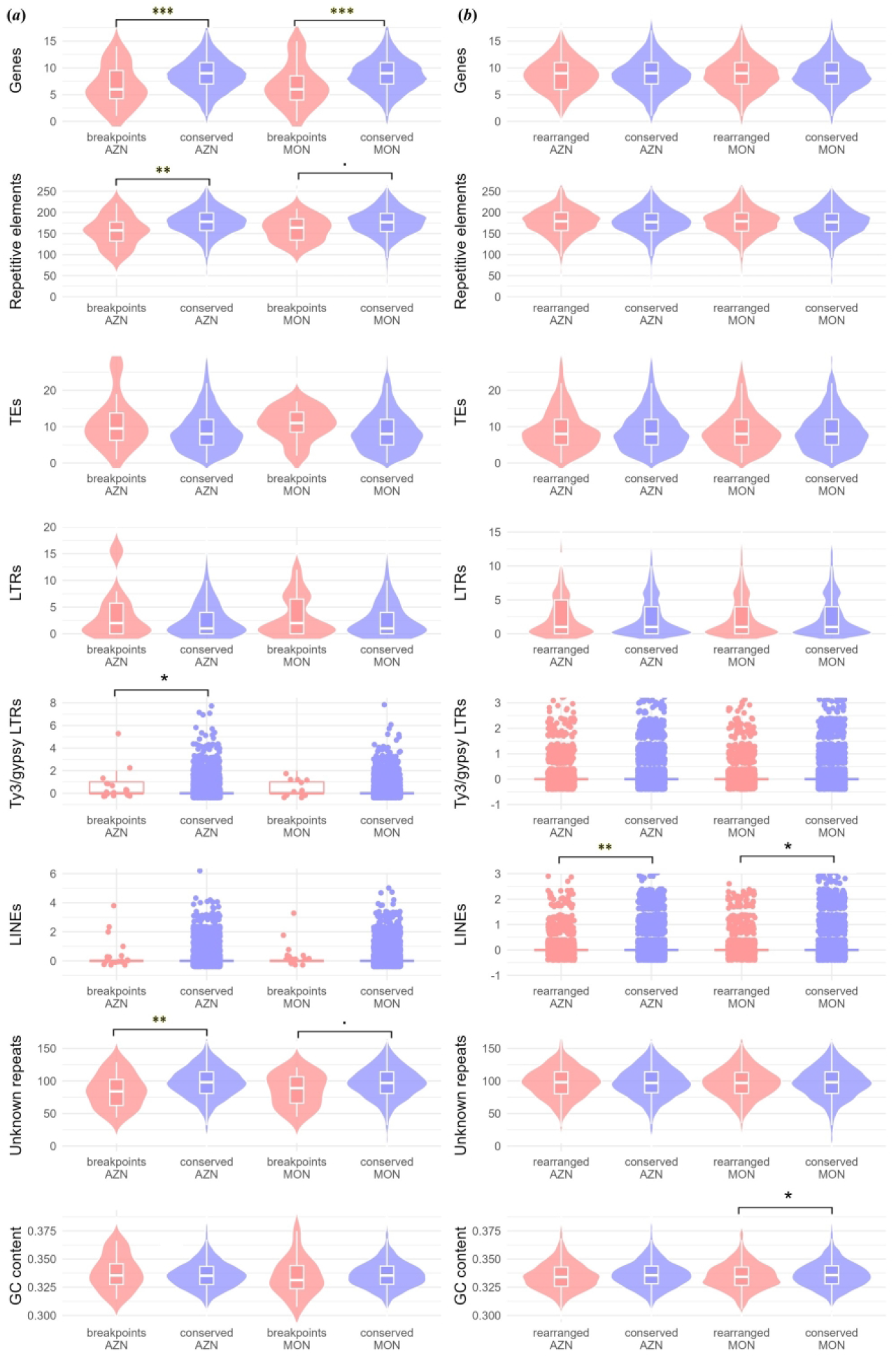
Comparison of genomic feature distributions between breakpoint and non-breakpoint regions, and across rearranged and collinear chromosomes. Violin and jitter plots showing the distribution of genomic features in breakpoint regions versus the rest of the genome (*a*), and in rearranged versus conserved chromosomes (*b*), using a 100kb window distribution. Statistical significance is indicated with asterisks (. p < 0.1; *p < 0.05; **p < 0.01; ***p < 0.001), corresponding to the linear model comparisons. For each genomic variable, the number of elements per window is represented, except for GC content for which the proportion is represented. Abbreviations as follows: RE: Repetitive elements, TEs: Transposable Elements, LTRs: Long Terminal Repeat Retrotransposons and LINEs: Long Interspersed Nuclear Repeats.

Breakpoints similarly differed from the rest of the genome (figure 2*a*, table 1, electronic supplementary material, table S4-S5, figure S5) even when compared within the same sections (interstitial and subtelomeric) in the same chromosome (table 1). However, the magnitude of the differences varied between genomes, analysis and the breakpoint distance to the chromosome center. Gene density for interstitial breakpoints was almost half compared to similar regions, but breakpoints gene impoverishment was not notable within subtelomeric sections (figure 2, table 1, see electronic supplementary materials, table S4). TE enrichment near breakpoints was significant both within interstitial and subtelomeric sections for the AZN genome, mainly because of LTR enrichment, particularly those from the Ty3/gypsy subclass (figure 2, table 1, see electronic supplementary materials, table S4-5). TE enrichment was less clear for the MON genome, where the effect was marginally significant (i.e. 0.1> P >0.05) and only within interstitial areas and was not detectable according to the linear model analysis (see electronic supplementary materials, table S4). The most common TRC was enriched in breakpoints located in the subtelomeric section for the MON genome, which include breakpoints involved in fusion/fission. Unknown repeats, the main component of repetitive elements (see electronic supplementary materials, figure S6), are scarcer near breakpoints in theGLMs but not the permutation analyses where their effects were only suggestive for only the AZN genome (table 1, see electronic supplementary materials, table S4-5). GC content varied in the breakpoint regions, with the direction of the effect differing between genomes, sections and analysis, likely because of correlations with other variables such as with gene density and unknown repeats (see electronic supplementary materials, figure S6).

**Table 1:** Enrichment permutation test by genomic region for the (a) MON and (b) AZN genomes. Breakpoints are compared against the rest of the rearranged chromosome within the same genomic region. For each genomic feature, the table reports the z-score, observed value, 95% confidence intervals (CIs) fold enrichment and p-value.The following abbreviations have been used: REs: Repetitive elements, TEs: Transposable Elements, LTRs: Long Terminal Repeat Retrotransposons, DNA trans: DNA transposons; LINEs: Long Interspersed Nuclear Elements; SINE: Short Interspersed Nuclear Elements and TRC: Tandem Repeat Cluster.

| a) MON genome |  |  |  |  |  |  |
| --- | --- | --- | --- | --- | --- | --- |
| Genomic feature | Region | z-value | Observed | 95% CIs | Fold Enrichment | p-value |
| Genes | Interstitial | -2.27 | 22.00 | [25, 50] | 0.61 | 0.008 |
|  | Subtelomeric | -0.29 | 10.00 | [5, 19] | 0.91 | 0.481 |
| REs | Interstitial | -0.42 | 686.00 | [590, 822] | 0.96 | 0.343 |
|  | Subtelomeric | -1.10 | 177.00 | [142, 273.02] | 0.83 | 0.133 |
| TEs | Interstitial | 1.51 | 53.00 | [17, 60.02] | 1.47 | 0.081 |
|  | Subtelomeric | -0.15 | 10.00 | [2, 24] | 1.00 | 0.518 |
| LTRs | Interstitial | 1.53 | 21.00 | [1, 25] | 2.10 | 0.091 |
|  | Subtelomeric | 0.17 | 4.00 | [0, 12] | 1.67 | 0.390 |
| Ty1/copia | Interstitial | 0.98 | 6.00 | [0, 9] | 1.50 | 0.213 |
|  | Subtelomeric | -0.18 | 1.00 | [0, 4] | 1.00 | 0.660 |
| Ty3/gypsy | Interstitial | 0.54 | 2.00 | [0, 5] | 1.50 | 0.367 |
|  | Subtelomeric | -0.53 | 0.00 | [0, 2] | 1.00 | 0.723 |
| DNA transp | Interstitial | 0.88 | 3.00 | [0, 5] | 1.50 | 0.259 |
|  | Subtelomeric | 0.70 | 1.00 | [0, 2] | 2.00 | 0.401 |
| LINEs | Interstitial | -0.85 | 0.00 | [0, 4] | 0.50 | 0.444 |
|  | Subtelomeric | -0.49 | 0.00 | [0, 2] | 1.00 | 0.768 |
| SINEs | Interstitial | -0.45 | 0.00 | [0, 2] | 1.00 | 0.760 |
|  | Subtelomeric | -0.27 | 0.00 | [0, 1] | 1.00 | 0.918 |
| Unknown | Interstitial | -0.19 | 385.00 | [305.97, 478] | 0.98 | 0.432 |
|  | Subtelomeric | -0.82 | 98.00 | [72.97, 163.02] | 0.84 | 0.215 |
| TRC | Interstitial | -0.65 | 0.00 | [0, 3] | 1.00 | 0.624 |
|  | Subtelomeric | 2.05 | 1.00 | [0, 1] | 2.00 | 0.115 |
| Most common TRC | Interstitial | -0.55 | 0.00 | [0, 2] | 1.00 | 0.710 |
|  | Subtelomeric | 2.65 | 1.00 | [0, 1] | 2.00 | 0.086 |
| GC content | Interstitial | -1.37 | 0.33 | [0.32, 0.35] | 0.98 | 0.083 |
|  | Subtelomeric | 0.84 | 0.34 | [0.32, 0.35] | 1.02 | 0.192 |

b) AZN genome
| Genomic feature | Region | z-value | Observed | 95% CIs | Fold Enrichment | p-value |
| --- | --- | --- | --- | --- | --- | --- |
| Genes | Interstitial | -3.45 | 17.00 | [28, 54] | 0.42 | 0.00 |
|  | Subtelomeric | 0.16 | 4.00 | [0, 8] | 1.33 | 0.49 |
| REs | Interstitial | -1.46 | 690.00 | [657.95, 900.02] | 0.88 | 0.09 |
|  | Subtelomeric | -1.56 | 40.00 | [28.98, 103.02] | 0.57 | 0.06 |
| TEs | Interstitial | 2.79 | 73.00 | [20, 64] | 1.82 | 0.00 |
|  | Subtelomeric | 2.42 | 12.00 | [0, 12] | 4.00 | 0.03 |
| LTRs | Interstitial | 2.83 | 31.00 | [2, 28.02] | 2.82 | 0.01 |
|  | Subtelomeric | 3.45 | 8.00 | [0, 7] | 4.27 | 0.01 |
| Ty1/copia | Interstitial | 1.29 | 7.00 | [1, 9] | 1.75 | 0.16 |
|  | Subtelomeric | 2.43 | 2.00 | [0, 2] | 2.15 | 0.07 |
| Ty3/gypsy | Interstitial | 3.16 | 6.00 | [0, 5] | 2.91 | 0.02 |
|  | Subtelomeric | 2.55 | 1.00 | [0, 1] | 1.82 | 0.09 |
| DNA transp | Interstitial | -0.05 | 2.00 | [0, 6] | 1.00 | 0.67 |
|  | Subtelomeric | -0.40 | 0.00 | [0, 1] | 0.86 | 0.85 |
| LINEs | Interstitial | -0.19 | 1.00 | [0, 5] | 0.89 | 0.63 |
|  | Subtelomeric | -0.30 | 0.00 | [0, 2] | 0.89 | 0.91 |
| SINEs | Interstitial | -0.46 | 0.00 | [0, 2] | 0.73 | 0.73 |
|  | Subtelomeric | -0.17 | 0.00 | [0, 1] | 0.97 | 0.97 |
| Unknown | Interstitial | -0.96 | 388.00 | [344, 512.05] | 0.90 | 0.17 |
|  | Subtelomeric | -1.74 | 17.00 | [15.98, 66] | 0.44 | 0.05 |
| TRC | Interstitial | -0.59 | 0.00 | [0, 2] | 0.69 | 0.68 |
|  | Subtelomeric | -0.20 | 0.00 | [0, 1] | 0.95 | 0.96 |
| Most common TRC | Interstitial | -0.57 | 0.00 | [0, 2] | 0.71 | 0.70 |
|  | Subtelomeric | -0.18 | 0.00 | [0, 1] | 0.96 | 0.97 |
| GC content | Interstitial | 1.50 | 0.35 | [0.32, 0.35] | 1.03 | 0.08 |
|  | Subtelomeric | 4.32 | 0.44 | [0.3, 0.39] | 1.30 | 0.00 |

### (e) Divergence and evolution in rearrangements

Substitution rates and selection pressure (ω = d_N_ / d_S_) was calculated for the 6 (out of 8 rearrangements) that reached the minimum number of sites required for the analysis following strict coding sequence (CDS) filtering (see electronic supplementary material, table S3). Two inversions (Inv_chr6 and Inv_chr32) showed marginally significant signs of altered gene evolution (table 2) but in opposite directions. Inv_chr6 showed patterns consistent with purifying selection (ω = 0; P = 0.059) with no non-synonymous mutations inside the inversion (d_N_ = 0; P = 0.050). In contrast, Inv_chr32 showed higher divergence both for synonymous (d_S_ = 0.29; d_S_ P = 0.059) and non-synonymous mutation rate (d_N_ = 0.66; P = 0.050) but with no apparent effect on selection efficacy (ω).

**Table 2:** Gene evolution permutation test inside areas affected by chromosomal rearrangements. Synonymous (dS), nonsynonymous (dN) substitution rates and their ratio (ω = dN/dS) are calculated through aggregated analyses, were substitutions and sites were summed across coding sequences (CDS) within each rearrangement and compared with areas of the same size permuted across the chromosome of the rearrangement.

| Rearran-<br>gements | Observed<br>(dN) | Observed<br>(dS) | Observed<br>($\omega$) | Predicted<br>(dN) | Predicted<br>(dS) | Predicted<br>($\omega$) | 95% CI<br>(dN) | 95% CI<br>(dS) | 95% CI<br>($\omega$) | p-value<br>(dN) | p-value<br>(dS) | p-value<br>($\omega$) |
| --- | --- | --- | --- | --- | --- | --- | --- | --- | --- | --- | --- | --- |
| Inv_chr8 | 0.12 | 0.11 | 1.10 | 0.14 | 0.14 | 0.97 | [0.03, 0.23] | [0.03, 0.24] | [0.38, 3.19] | 0.287 | 0.277 | 0.208 |
| Inv_chr6 | 0.00 | 0.01 | 0.01 | 0.10 | 0.05 | 0.85 | [0, 0.77] | [0, 0.93] | [0, 3.49] | 0.050 | 0.109 | 0.059 |
| Inv_chr32 | 0.66 | 0.29 | 2.31 | 0.13 | 0.11 | 1.04 | [0, 0.66] | [0.01, 0.29] | [0.1, 4.02] | 0.050 | 0.059 | 0.327 |
| Trans_chr16 | 0.04 | 0.04 | 0.94 | 0.19 | 0.10 | 0.99 | [0, 1.66] | [0, 1.17] | [0.06, 5.14] | 0.228 | 0.366 | 0.475 |
| Trans_chr24 | 0.01 | 0.02 | 0.70 | 0.01 | 0.09 | 0.54 | [0, 0.62] | [0, 0.63] | [0, 2.04] | 0.436 | 0.317 | 0.683 |
| Trans_chr5 | 0.03 | 0.05 | 0.51 | 0.03 | 0.10 | 0.68 | [0, 0.76] | [0, 0.64] | [0.21, 1.61] | 0.436 | 0.337 | 0.356 |

### (f) Gene ontology across rearranged regions

The eggNOG-mapper successfully assigned functional orthology to 23,975 genes for MON (76.8% of the total annotated genes) with 453 GO annotated genes in regions affected by CRs (70.8% of all annotated genes within the CRs): 39 in Inv_chr 6 (65%), 184 in Inv_chr8 (68.4%), 83 in Inv_chr32 (85.6%), 34 in Trans_chr5 (85.5%), 24 in Trans_chr9 (88.9%), 21 in Trans_chr16 (56.8 %), 40 in Trans_chr24 (67.8%), 9 in Fus_chr17 (69.2%) and 19 in Fus_chr19 (65.5%).

Gene Ontology enrichment analyses for all CRs combined and separately for each CR showed that rearranged regions are enriched for a wide variety of cellular functions, with 188 enriched GOs for biological processes (BP), 51 for molecular functions (MF) and 47 for cellular compartments (CC) (see electronic supplementary material, table S6). These GO terms are mainly focused on transcription regulation, protein glycosylation, proteasome activation and DNA topoisomerase (Figure 3; see electronic supplementary material, table S6). These functions are enriched mainly in Inv_chr6, Inv_chr8 and Trans_chr16. However, other CRs are specialised in other functions: Trans_chr5 regulate carbohydrate catabolism, Trans_chr9 manage chloroplast transmembrane transport and oxidoreductase activity and Trans_chr24 covers acquired resistance and Inv_chr32 is focused on response to cold and phosphate starvation (see electronic supplementary material, table S6).

**Figure 3:**
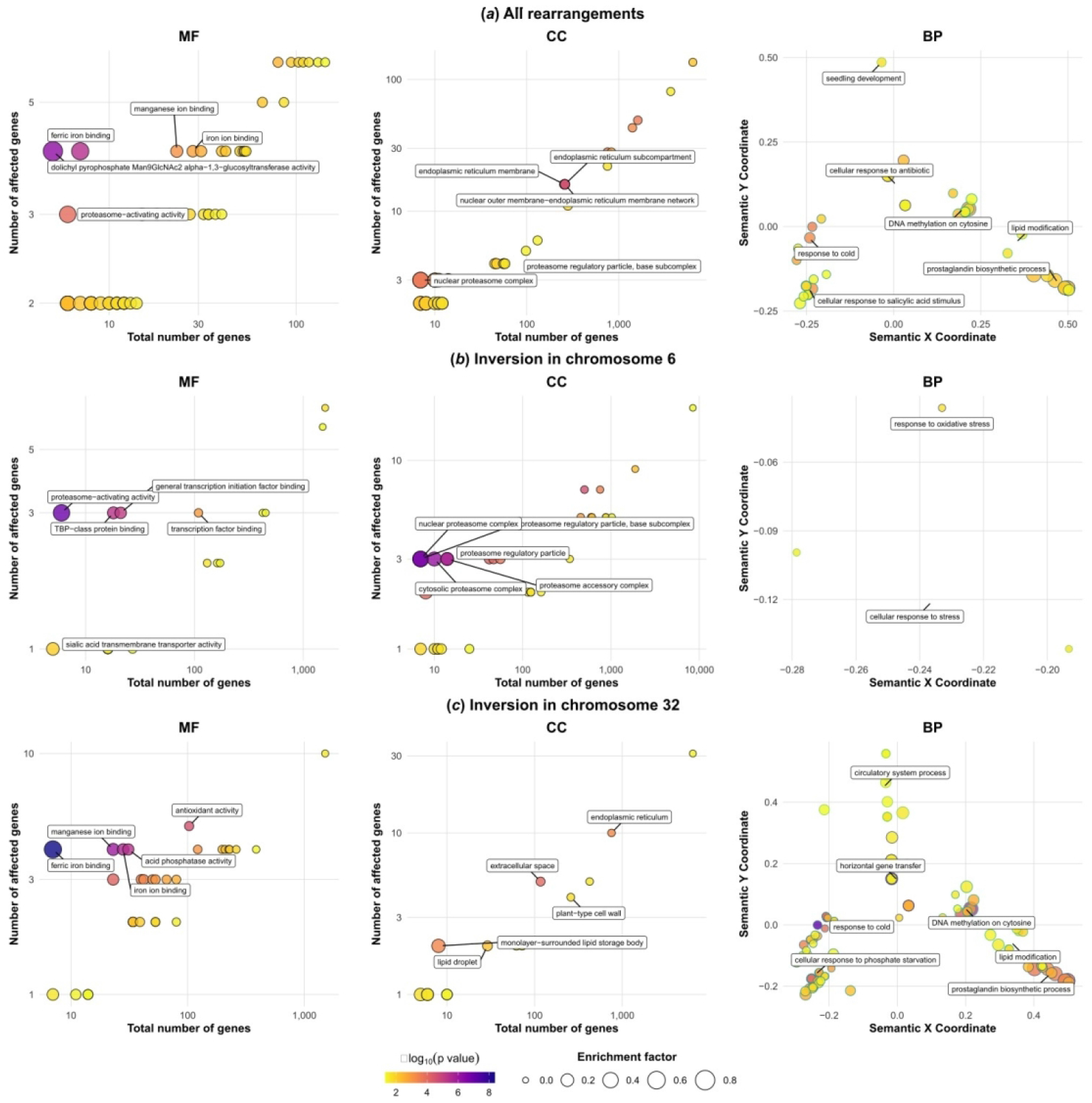
Functional enrichment and semantic clustering of Gene Ontology (GO) terms for genomic rearrangements. Semantic clustering and enrichment of Gene Ontology (GO) terms for all rearrangements (*a*), inversion in chr 6 (*b*) and chr 32 (*c*) for MON. The three function types are represented from left to right: Molecular Function (MF), Cellular Component (CC) and Biological Process (BP). Molecular Function and CC are represented through scatter plots where the x-axis represents the number of annotated genes throughout the genome and y-axis the number of genes in the area affected by the rearrangement. BP is represented through semantic plots, visualizing it through the 2D semantic space via Multidimensional Scaling (MDS). Highly redundant terms were grouped using GOSemSim (Resnik measure based on *Arabidopsis thaliana* annotations) and were labeled by the most representative term. In all the plots, point size indicates enrichment factor and color represents –log₁₀(p-value).

## Discussion

### (a) A colonization through founder events reveals a striking number of rearrangements

West-to-east decreasing dysploidy across European *C. helodes* populations was reported by Escudero and colleagues [60]. Using genomic tools, eight CRs (1 fusion, 3 inversions and 4 translocations) were detected between individuals from the two geographically extreme populations of the species’ European range (figure 1), a much higher number than expected based on traditional cytogenetic analyses (figure 1; [61]). The number of CRs is striking considering the recent history of this species, as the estimated age of the crown node for the European populations of *C. helodes* is 40,000 years [61]. This would translate to at least one CR per 10,000 years [60] which is much higher than any previous estimations for angiosperms [100], even considering other species of the karyologically dynamic genus *Carex* [54,56,101].

Consistent with former inferences [60], the identified CRs could have been established through recurrent genetic bottlenecks during range expansion. This hypothesis is further supported by the fact that the assembled genomes show the only (MON) or the most common (AZN) euploid karyotype for each population. Accordingly, fusions and other cytologically detectable CRs between these two euploid chromosome numbers (2n = 36^II^ and 2n = 35^II^) are less likely to occur within each population (figure 1*b*; [60]). During range expansion, even underdominant CRs can establish and proliferate [1,18]. A long life cycle [61] coupled with the possibility for asexual reproduction further promotes the establishment and persistence of underdominant CRs. Selfing capability, evidenced by our crossing experiment and the high homozygosity in natural populations [61] provide a plausible mechanism for this hypothesis: individuals with new CRs may produce seeds by selfing that colonize and establish in new areas [102]. However, our hypothesis for CRs that are not detectable by cytology, such as inversions or translocations, remains tentative until the fixed or polymorphic status of these CRs is confirmed in each population through population-level genomic data. Indeed, one inversion was heterozygotic for MON. Nevertheless, this conclusion remains consistent for CRs that are detectable through cytology such as chromosome fissions and fusions [61].

Hybrid dysfunction led by CR accumulation has been reported in other *Carex* species [103,104]. While there is no strong reproductive isolation between our studied populations as revealed by F1s high germination success, reproductive isolation increases in the following generation as F2s reach a germination success of only 60% (significantly lower than in natural populations [109]). This increased hybrid dysfunction could be potentially explained by partial sterility of the F_1_s due to chromosome missegregation during meiosis. In congruence with this, hybrid dysfunction was already reported in *C. scoparia* where germination success decreases as chromosome differences increase between parents [103].

However, multi-generational fitness measurements and more exhaustive cytogenetic studies would be necessary to clarify the extent and impact of hybrid dysfunction. The accumulation of CRs preceding genetic differentiation [60,62] may suggest that CRs are more likely to be drivers of genetic differentiation rather than a consequence of it.

### (b) Gene-poor and TE-rich environments in the genomic landscape facilitate viable CRs

Both breakpoints and rearranged chromosomes showed distinct genomic characteristics from the rest of the genome. This is consistent with previous findings suggesting that rearrangements are restricted and constrained to genomic regions with particular genomic features [25,31–33], including *Carex* [25] and other holocentric organisms [26]. The observed reduced gene density near breakpoints compared to the rest of the genome (table 1) support the hypothesis that CR breakpoints lie in gene poor areas which reduce the potential deleteriousness of CRs [25,33], which has been observed for both mono [33,106] and holocentric organisms [25,26,107].

Transposable elements (TEs) were overall enriched near the breakpoints yet the magnitude of the enrichment depended on the analysed genome (table 1), within TEs the most enriched were LTRs, especially Ty3/Gypsy subclass (table 1, figure 2, see electronic supplementary material, table S4-S5). TEs enrichment appears to be more local than gene impoverishment as its effect was less notable in the GLM analysis (figure 2, see electronic supplementary material, table S4). The association between TEs and LTRs with CRs has been consistently reported across different organisms [25,26,112,113]. Evidence at both macro- [109] and microevolutionary (shown here) scales support the role of LINEs and LTR (Ty3/Gypsy) in chromosome instability, as rearranged chromosomes tend to be enriched in these repetitive elements. TEs can cause CRs through non-allelic homologous recombination (NAHR) or alternative transposition for DNA transposons [110,111]. However, CRs can also lead to TE accumulation through recombination suppression [116,118,119]. In addition, TE rich regions are also gene-poor [47,48], so the observed correlation could also reflect a higher likelihood of CRs to establish due to their lower deleteriousness. The available data does not allow us to disentangle which of these explanations for the association is responsible for the observed pattern.

In many monocentric species, gene density and GC content decrease toward the centromere, increase toward the telomeres, and subsequently decline sharply at chromosomal ends, exhibiting an inverse relationship with repeat content [47–49]. Other patterns have been reported in holocentric organisms [43–46]. The uniformity of genomic characteristics for holocentric plants was only suggested for gene density in our data as distance to the chromosome center is an important predictor for nearly all the genomic characteristics measured (see electronic supplementary material; figure S4, table S4).

### (c) Contrasting evolutionary trajectories and functional signatures of chromosomal inversions

Among the observed CRs, the inversions on chromosomes 6 and 32 were the only ones with distinct gene evolution patterns compared to the rest of the chromosome (table 2), but in opposite directions. The inversion on chromosome 32 showed increased neutral and non-synonymous divergence and, therefore, it did not translate into difference in selection pressure (ω). Such increased divergence is consistent with a role of inversions as islands of genetic differentiation, likely due to recombination suppression and linkage disequilibrium within heterozygotic karyotypes [114]. Notably, this CR is the only one with heightened neutral divergence, pointing to it being likely older than the other CRs, further supporting the hypothesis of its appearance prior to the Spanish colonization as our Portuguese individual was heterozygous for this CR (see electronic supplementary materials, table S2, figure S3). In contrast, the inversion on chromosome 6 showed patterns consistent with purifying selection (table 2). This typically points to important functions being located within these inversion, so mutations are potentially deleterious for the plant. While the inversion capturing important genes by chance is perfectly possible, purifying selection can be detected in non-fundamental genes when they are in linkage desequilibrium with fundamental ones, as it happens with genes within an inversion [115,116].

Our GO enrichment analysis offers an exploratory insight into the potential functional implications of the identified inversions. Therefore, these results must be interpreted with caution due to the inherent limitations of cross-species GO transfer and incomplete genome annotations. Inv_chr6 is enriched in functions related to positive regulation of proteasome activity, which has been linked to increased tolerance for CRs [117,118]. Therefore it could mitigate proteotoxic stress arising from gene dosage imbalances that may come from duplicated and/or eliminated parts in translocations [119–121]. Notably, nuclear proteasomes can also increase CR rate, as they degrade trapped topoisomerases exposing double strand breaks (DSBs) for repair [122,123], but also promote their repair through CR-prone mechanisms [124,125].

The inversion on chromosome 32 (Inv_chr32) is enriched for functions related to cold response and phosphate starvation, matching the geological and climatic contrast between both localities: Aznalcóllar features hot, dry summers and is located in the the Iberian Pyrite Belt with acidic, sulfurous soils which limits phosphorus availability [126]. Conversely, the mountainous Monchique is a high-precipitation refuge with milder temperatures and less sulfurous soils [127]. This preliminary finding opens the door for future experimental studies to confirm and explore the potential role of these inversions in population differentiation and local adaptation.

## Final Remarks

The synteny analysis characterized a striking number of CRs between the two typical karyotypic races of our *C. helodes* populations, suggesting a high rearrangement rate even for the already kariologically dynamic genus *Carex*. CRs establishment was likely facilitated by genetic bottlenecks during the colonization of southwestern Spain. Breakpoints were gene-poor and TE-rich, characteristics that facilitate the occurrence and/or limit the CRs deleteriousness, likely accounting for some of the high number of CRs. Rearranged chromosomes as a whole also have distinct genomic characteristics, such as an increased LINE content, supporting their possible role in CRs as was previously reported for *Carex* [109]. Two inversions showed suggestive signs of distinct gene evolution in opposite directions. The inversion on chromosome 6 shows a pattern that is consistent with early signs of purifying selection, aligning with its enrichment in proteasomal activity, related to CR tolerance. The inversion on chromosome 32 shows increased divergence between the two genomes and was enriched in functions related to response to temperature stress and phosphorus deficit, two environmental factors that differ between the two localities.

Ultimately, the accumulation of CRs and breakpoints with specific genomic features, suggest rapid intraspecific karyotypic evolution in *C. helodes*. These mechanisms likely serve as the foundational architecture for eventual differentiation, where chromosomal barriers effectively sequester potential adaptive loci facilitating differential genomic differentiation.

## Acknowledgements

The authors wish to thank Ms Nafiseh Sargheini for helpìng and teaching with Hi-C extraction and Ms Alegria Montero for performing DNA extraction, Dr Ignacio Márquez- Corro for helping with field work and caring for the plants in the greenhouse and Mr Diego Herrero Doblado continuing greenhouse work, performing the experimental crosses and estimating germination success.

## Funding information

This work was funded by the Spanish Government and FEDER funds (EuropeanCommission) through projects PID2021-122715NB-I00 and PID2024-157198NB-I00 and by the Andalusian regional government (Spain) through project ProyExcel_00125. The latter also supported SM-B and RS-V. IG-R was supported by the Andalusian Regional Ministry of Economy, Knowledge, Business, and University (PREDOC_00632). KL was supported by the Swiss National Science Foundation grants PCEFP3_202869 and 220868 as well as the Fondation Pierre Mercier pour la Science and the Fondation du Jardin Botanique de Neuchâtel.

## Notes

### Competing Interest Statement

The authors have declared no competing interest.

