## Supplementary tables and figures captions for "Genomic insights into the karyotypic radiation of a narrow endemic holocentric plant *Carex helodes*"

**Figures supplementary materials:**

**Figure S1: Telomeric repeat identification (TIDK) of all preliminary pseudomolecules of MON (*a*) and AZN (*b*) genome.**

Visualization of the sequence composition across all 36 and 35 preliminary pseudomolecules for MON (*a*) and AZN (*b*) genome, respectively.

**Figure S2: Reciprocal Hi-C cross-mapping contact maps.**

Physical validation of computationally derived chromosomal rearrangements. The contact map reveals characteristic off-diagonal "bowtie" interaction signatures at the exact genomic coordinates of inversions. For interchromosomal CRs (fusion, fission and translocations), different domains are clearly visible in the chromosomes involved, demonstrating that they were in different chromosomes in the other genome.

**Figure S3: Hi-C contact map for chromosomes with inversions for MON (*a*) and AZN (*b*) genome**

Physical validation of the three inversions heterozygosity for MON (*a*) and AZN (*b*) genome, respectively. As the Hi-C reads are aligned against their own genome, inversions characteristic off-diagonal "bowtie" interaction signatures are only expected for heterozygous inversions.

**Figure S4: Genomic features along all chromosomes.**

Distribution of genes, transposable elements (TEs) and sequence features along

all chromosomes for MON (a) and AZN genome (b). Each panel shows observed

values (grey dots) and smoothed trends (colored lines) across genomic windows for genes,

total TEs, LTRs, Ty1/copia, Ty3/gypsy, LINEs, unclassified repeats, tandem repeats, and GC

content. Vertical grey lines indicate rearrangement breakpoints.

**Figure S5: Genomic features along rearranged chromosomes.**

Distribution of genes, transposable elements (TEs) and sequence features along

rearranged chromosomes for MON (a) and AZN genome (b). Each panel shows observed

values (grey dots) and smoothed trends (colored lines) across genomic windows for genes,

total TEs, LTRs, Ty1/copia, Ty3/gypsy, LINEs, unclassified repeats, tandem repeats, and GC

content. Vertical colored lines indicate rearrangement breakpoints and their correspondence

between the two genomes.

**Figure S6: Correlation matrix of genomic features for Carex helodes genomes**

Lower triangular matrix showing the relationship between different genomic attributes across 100-kb windows for MON (*a*) and AZN genome (*b*). Features are ordered by hierarchical clustering. The colors represent the correlation sign: negative correlations are represented in blue and positive ones in blue. The intensity of the color corresponds to the Pearson correlation coefficient. The following abbreviations have been used: TEs: Transposable Elements, LTR: Long Terminal Repeat Retrotransposons, DNA trans: DNA transposons; LINE: Long Interspersed Nuclear Repeats; SINE: Short Interspersed Nuclear Repeats, TRC: Tandem Repeat Cluster, genes: gene density and gc: GC content proportion.

**Table S1: BUSCO completeness assessment**

BUSCO completeness assessment for the genome assemblies and annotated gene sets for AZN and MON genome. Two BUSCO lineages were used as reference: Embryophyta and Poales.

**Table S2: Long-read split-read support for chromosomal rearrangements in both genomes.**

Genomic coordinates and sequencing read support for high-confidence breakends (BNDs) identified in the MON and AZN genome. Columns indicate the chromosome or scaffold name, exact genomic position (bp), variant identifier, and the assigned genotype (GT; 0/1 for heterozygous, 1/1 for homozygous alternative). Allelic support is broken down into reference allele depth and alternative allele depth, followed by the total depth of coverage at the locus. Variant calling was performed using *pbsv* with default parameters for structural variant detection.

**Table S3: Gene and CDS filtering statistics for evolutionary analysis.**

Number of genes/CDS (Coding DNA Sequences) inside rearranged areas and outside them within their rearranged chromosome in each step of the filtering to obtain adequate CDS to perform gene evolution analysis. N genes and N CDS refer to the total number of genes and CDS respectively within the annotated gene set respectively. Fifth column refers to the number of CDS with enough alignment (<10 % gaps and length above 120 bp) with good column alignment. Sixth column are CDS with adequate dN (non-synonymous substitution rates) and dS (synonymous substitution rates) values for gene evolution analysis.

**Table S4: Generalized linear models comparing genomic characteristics between breakpoints, rearranged and collinear chromosomes for both genomes.**

Generalized linear models exploring how different genomic characteristics vary through the two genomes through 100kb windows. Model predictors include rearranged chromosomes (CR), rearrangement breakpoints (BP:RC) and the distance to the chromosome midpoint (DC) and the interaction between them. Only results meeting significant (P < 0.05) or suggestive (P < 0.1) thresholds are shown.. Each sheet addresses one of the two genomes.

**Table S5: Permutation test for breakpoints enrichment in different genomic features for (*a*) MON and (*b*) AZN genomes**.

Breakpoints are compared against the rest of the rearranged chromosomes (excluding telomeres). For each genomic feature, the table reports the z-score, observed value, 95% confidence intervals (CIs), fold enrichment and p-value. The following abbreviations have been used: REs: Repetitive elements, TEs: Transposable Elements, LTRs: Long Terminal Repeat Retrotransposons, DNA trans: DNA transposons; LINEs: Long Interspersed Nuclear Repeats; SINEs: Short Interspersed Nuclear Repeats and TRC: Tandem Repeat Cluster.

**Table S6: Enriched gene ontology (GO) terms for all of the rearrangements, combined and each one separated.**

Enriched gene ontology (GO) terms for all of the rearrangements. combined and each one separated. All of the significant GO terms are shown for the three function types: BP (Biological Process), CC (Cellular Compartment) and MF (Molecular Function), according to the classic Fisher statistical test.

**Supplementary methods:**

Supplementary methods regarding: (*a*) Experimental crosses, germination rate estimation and cytogenetic study, (b) De novo repeat discovery and annotation and (c) CRs validation and inversion heterozygosity.
