## Supplementary figures and images for "Genomic insights into the karyotypic radiation of a narrow endemic holocentric plant *Carex helodes*"

### Supplementary figure S1

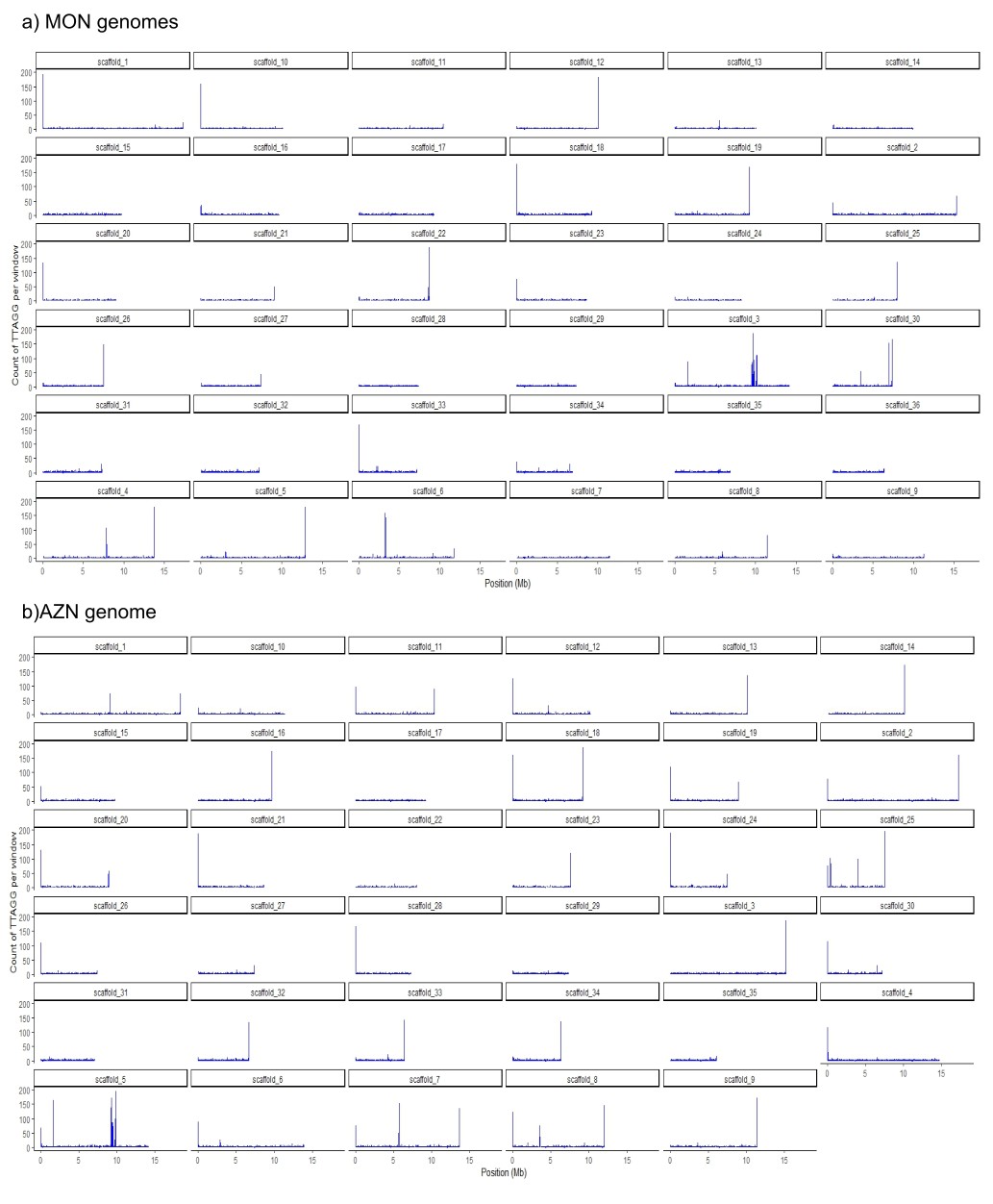

### Supplementary figure S2

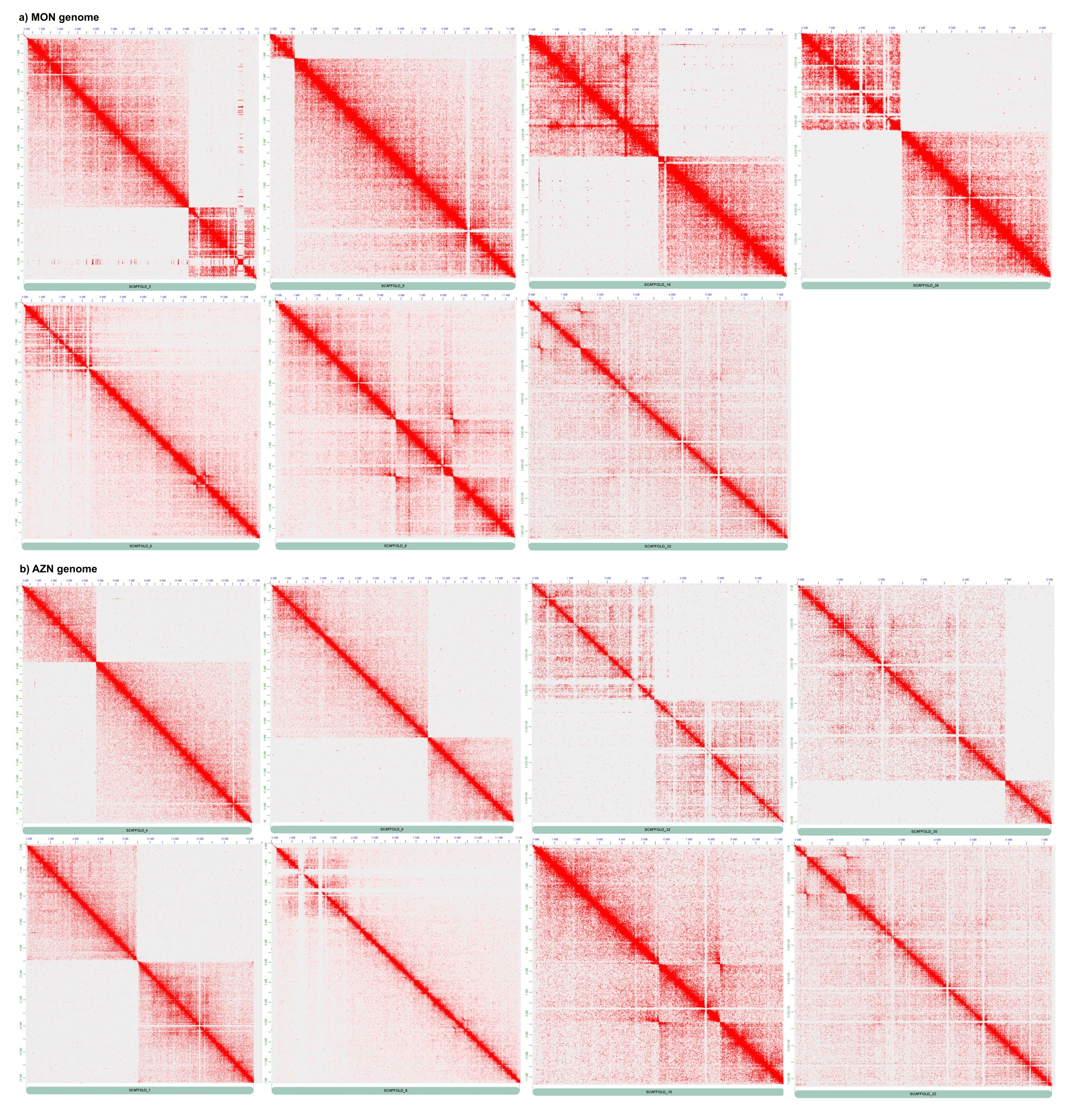

### Supplementary figure S3

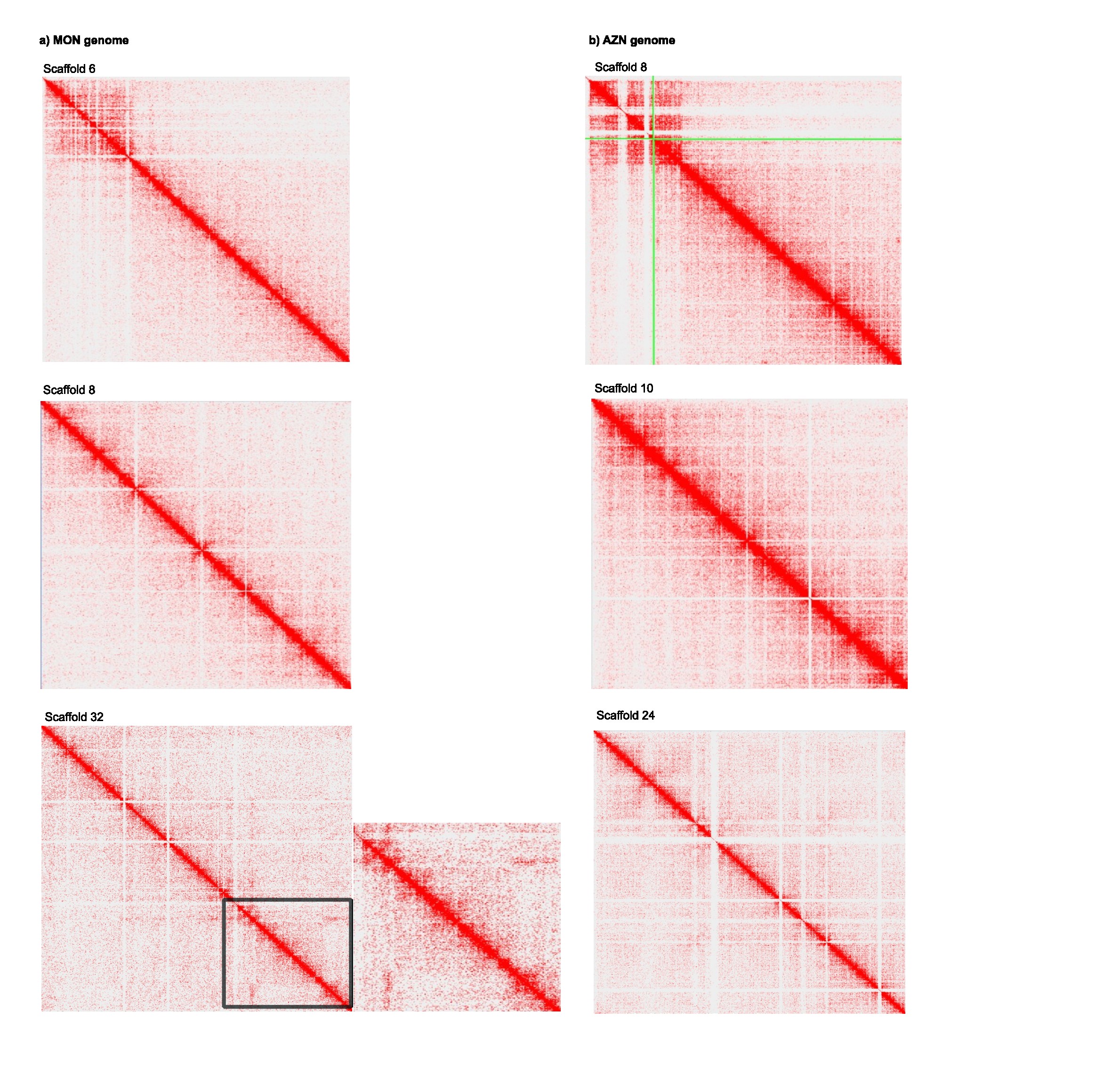

### Supplementary figure S4

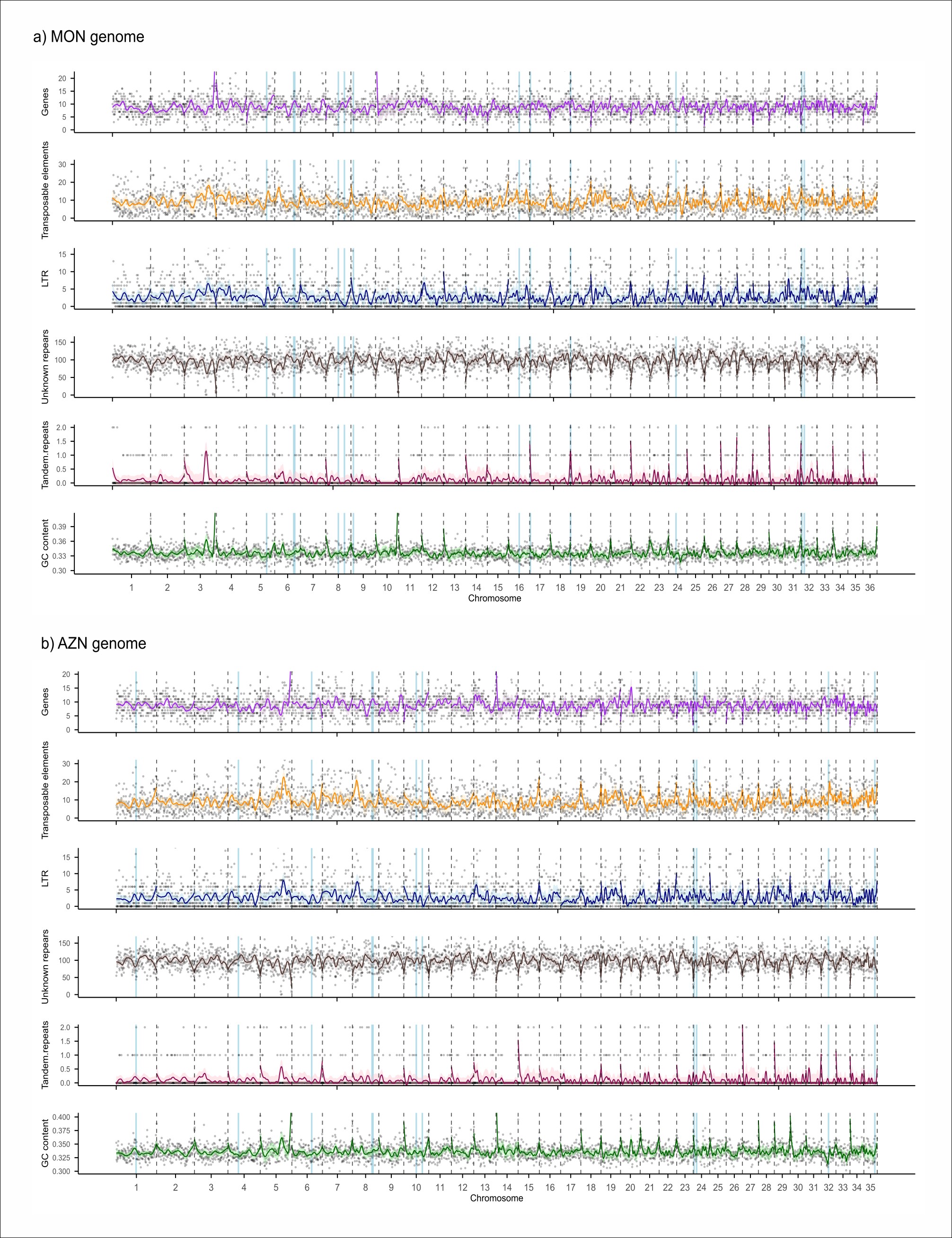

### Supplementary figure S5

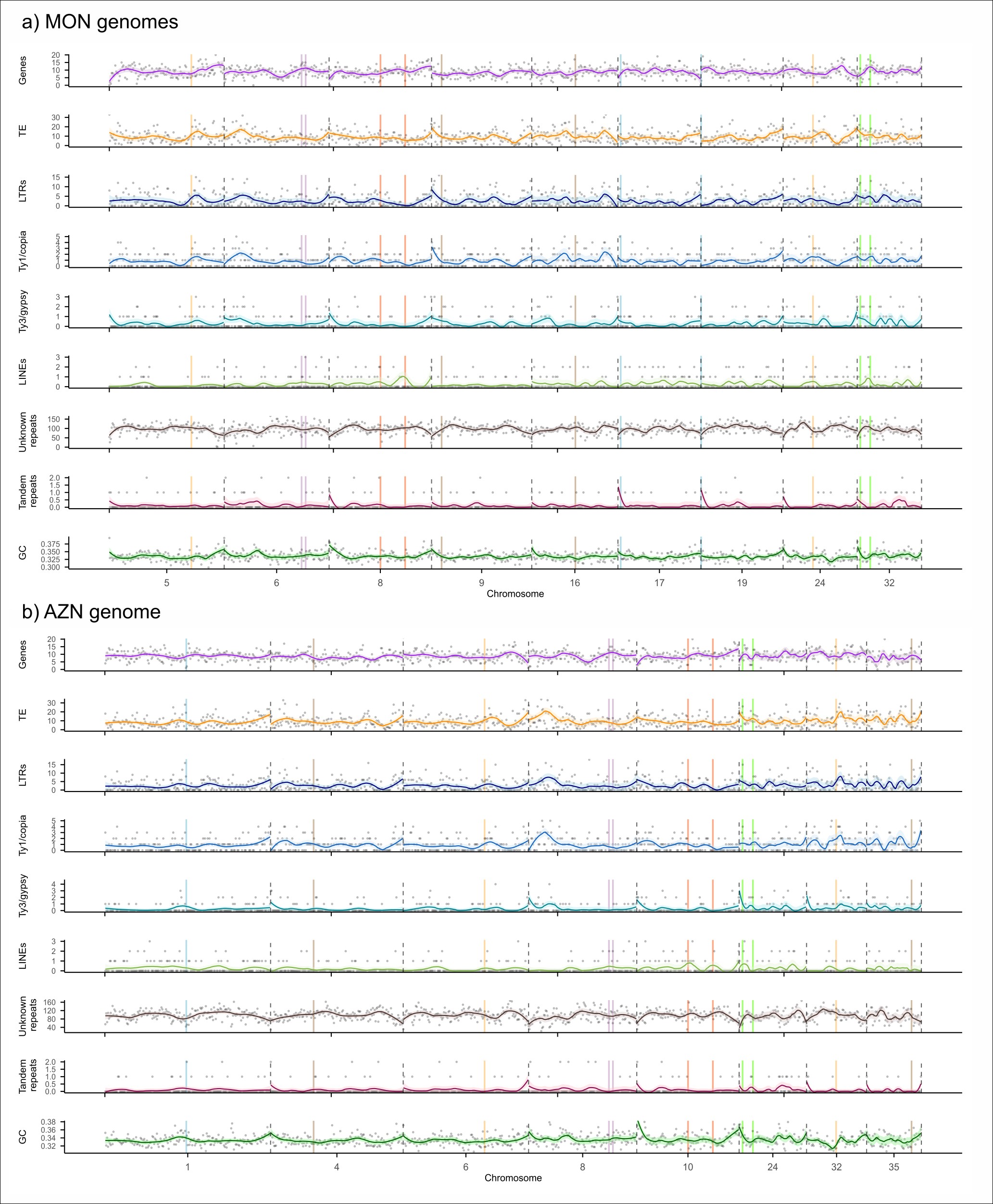

### Supplementary figure S6

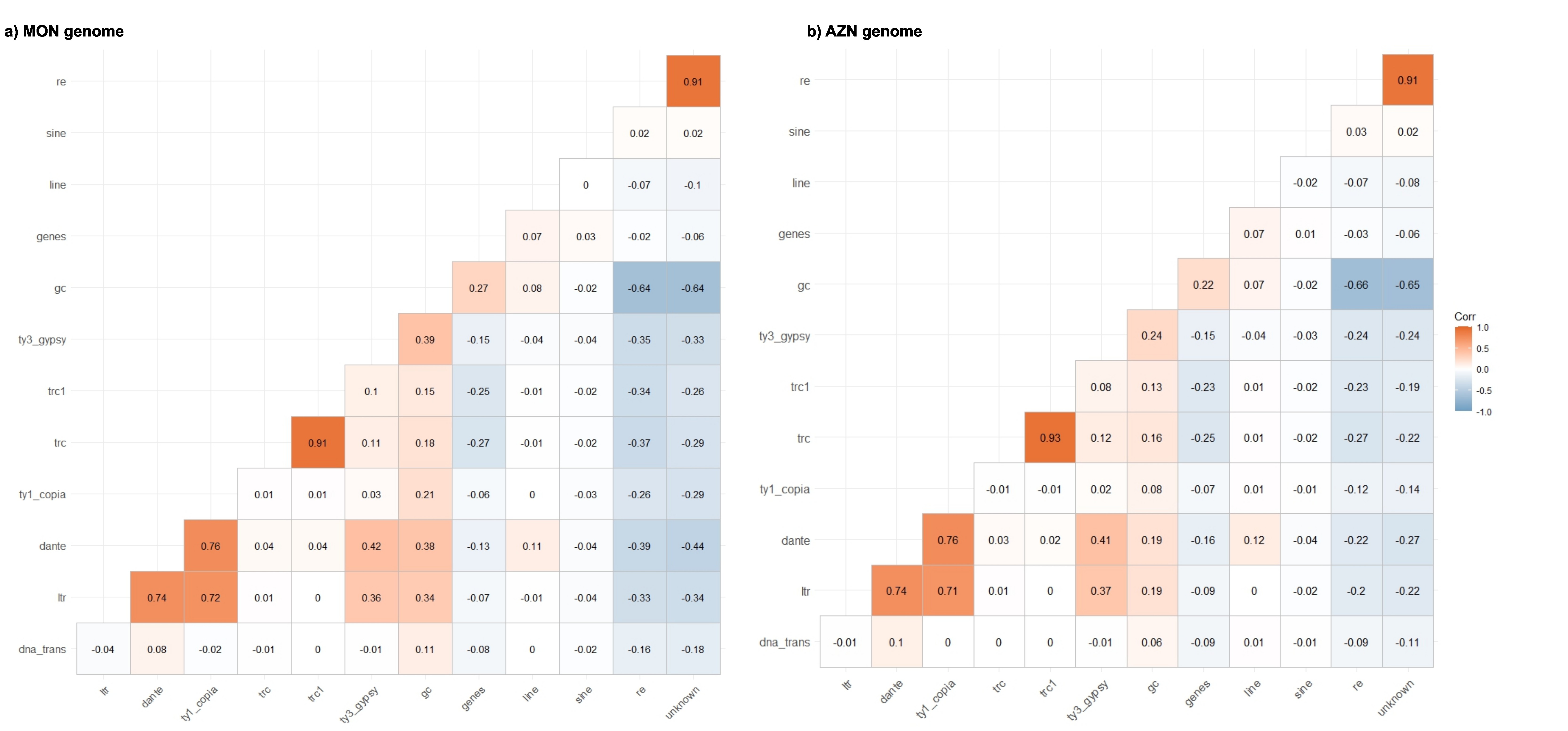
