## Supplementary table S1 for "Genomic insights into the karyotypic radiation of a narrow endemic holocentric plant *Carex helodes*"

**Table S1: BUSCO completeness assessmen**t: for the genome assemblies, for genome assembly and annotated gene sets for AZN and MON genome. Two BUSCO lineages were used as reference: Embryophyta and Poales.

|  | AZN | | | | |  | MON | | | | |
| --- | --- | --- | --- | --- | --- | --- | --- | --- | --- | --- | --- |
|  | Genome assembly | |  | Gene annotation | |  | Genome assembly | |  | Gene annotation | |
|  | Embryo-phyta | Poales |  | Embryo-  phyta | Poales |  | Embryo-  phyta | Poales |  | Embryo-  phyta | Poales |
| Single copy complete genes (S) | 93.68 % | 77.92 % |  | 78.5 % | 60.95% |  | 93.93% | 76.92 % |  | 77.63 % | 60.85 % |
| Duplicated complete genes (D) | 1.61 % | 3.84 % |  | 14.62 % | 20.53 % |  | 1.30 % | 3.84 % |  | 14.31 % | 19.89 % |
| Fragmented Genes (F) | 1.95 % | 0.98 % |  | 2.11 % | 0.88 % |  | 0.99 % | 0.98 % |  | 2.66 % | 0.88 % |
| Incomplete Genes (I) | 0 % | 0 % |  | 0 % | 0 % |  | 0 % | 0 % |  | 0 % | 0 % |
| Missing Genes (M) | 3.66 % | 18 % |  | 4.77 % | 17.65 % |  | 3.78 % | 18.22 % |  | 5.39 % | 18.30 % |
