## Supplementary table S3 for "Genomic insights into the karyotypic radiation of a narrow endemic holocentric plant *Carex helodes*"

Table S3: **Gene and CDS filtering statistics for evolutionary analysis.** Number of genes/CDS (Coding DNA Sequences) inside rearranged areas and outside them within their rearranged chromosome in each step of the filtering to obtain adequate CDS to perform gene evolution analysis. N genes and N CDS refer to the total number of genes and CDS respectively within the annotated gene set respectively. Fifth column refers to the number of CDS with enough alignment (<10 % gaps and length above 120 bp) with good column alignment. Sixth column are CDS with adequate dN (non-synonymous substitution rates) and dS (synonymous substitution rates) values for gene evolution analysis.

| Rearrangement | In/Out | N genes | N CDS | N CDS after alignment | N CDS ready for gene evolution |
| --- | --- | --- | --- | --- | --- |
| Inv_chr6 | In | 60 | 268 | 13 | 5 |
|  | Out | 945 | 4578 | 1467 | 153 |
| Inv_chr8 | In | 259 | 1383 | 86 | 21 |
|  | Out | 737 | 3779 | 531 | 152 |
| Inv_chr32 | In | 94 | 452 | 17 | 7 |
|  | Out | 557 | 2663 | 796 | 81 |
| Trans_chr16 | bp | 36 | 139 | 62 | 7 |
|  | Out | 754 | 2985 | 574 | 80 |
| Trans_ chr24 | bp | 57 | 167 | 49 | 16 |
|  | Out | 683 | 2732 | 839 | 104 |
| Trans_ chr5 | bp | 49 | 190 | 79 | 19 |
|  | out | 1117 | 4830 | 1563 | 215 |
| Trans_ chr9 | bp | 25 | 105 | 16 | 2 |
|  | Out | 834 | 4167 | 419 | 42 |
| Fusion chr17- chr19 | Bp | 40 (13 chr17 + 27 chr19) | 225 (70 chr17 + 155 chr19) | 38 | 5 |
|  | Out | 1556 (797 chr17 + 762 chr19) | 7177 (3596 chr17 + 3581 chr19) | 2466 | 260 |
