## Supplementary table S5 for "Genomic insights into the karyotypic radiation of a narrow endemic holocentric plant *Carex helodes*"

**Table S5: Permutation test for breakpoints enrichment in different genomic features for (a) MON and (b) AZN genomes**.

Breakpoints are compared against the rest of the rearranged chromosomes (excluding telomeres). For each genomic feature, the table reports the z-score, observed value, 95% confidence intervals (CIs), fold enrichment and p-value. The following abbreviations have been used: REs: Repetitive elements, TEs: Transposable Elements, LTRs: Long Terminal Repeat Retrotransposons, DNA trans: DNA transposons; LINEs: Long Interspersed Nuclear Repeats; SINEs: Short Interspersed Nuclear Repeats and TRC: Tandem Repeat Cluster.

a) MON genome

| **Genomic Feature** | **z-value** | **Observed** | **95% CIs** | **Fold Enrichment** | **p-value** |
| --- | --- | --- | --- | --- | --- |
| Genes | -2.20 | 27.00 | [31, 63.02] | 0.62 | 0.006 |
| REs | -0.59 | 809.00 | [713.95, 970.02] | 0.95 | 0.269 |
| TEs | 0.93 | 55.00 | [22, 69.05] | 1.28 | 0.187 |
| LTRs | 1.00 | 21.00 | [2, 32] | 1.62 | 0.163 |
| Ty1/copia | 0.63 | 6.00 | [1, 9] | 1.50 | 0.313 |
| Ty3/gypsy | 0.30 | 2.00 | [0, 5] | 1.50 | 0.428 |
| DNA transp | 0.59 | 3.00 | [0, 5.02] | 1.50 | 0.352 |
| LINEs | -0.97 | 0.00 | [0, 5] | 0.50 | 0.351 |
| SINEs | -0.52 | 0.00 | [0, 2] | 1.00 | 0.690 |
| Unknown | -0.27 | 457.00 | [380, 565] | 0.97 | 0.403 |
| TRC | 0.48 | 1.00 | [0, 3] | 2.00 | 0.378 |
| Most common TRC | 0.68 | 1.00 | [0, 2.02] | 2.00 | 0.345 |
| GC content | 0.39 | 0.34 | [0.33, 0.35] | 1.01 | 0.345 |

b) AZN genome

| **Genomic feature** | **z-value** | **Observed** | **95% CIs** | **Fold Enrichment** | **p-value** |
| --- | --- | --- | --- | --- | --- |
| Genes | -3.32 | 17.00 | [28, 55] | 0.42 | 0.001 |
| RE | -1.42 | 690.00 | [650.98, 900.02] | 0.88 | 0.085 |
| TE | 2.79 | 73.00 | [20, 65] | 1.82 | 0.008 |
| LTR | 2.73 | 31.00 | [1.98, 28] | 2.58 | 0.015 |
| Ty1/copia | 1.22 | 7.00 | [1, 9] | 1.75 | 0.162 |
| Ty3/gypsy | 3.49 | 6.00 | [0, 5] | 3.06 | 0.010 |
| DNA transp | 0.06 | 2.00 | [0, 5] | 1.00 | 0.563 |
| LINE | -0.13 | 1.00 | [0, 5] | 0.92 | 0.662 |
| SINE | -0.44 | 0.00 | [0, 2] | 0.76 | 0.758 |
| Unknown | -1.03 | 388.00 | [355.97, 524.02] | 0.90 | 0.154 |
| TRC | -0.59 | 0.00 | [0, 3] | 0.69 | 0.678 |
| Most common TRC | -0.53 | 0.00 | [0, 2] | 0.72 | 0.728 |
| GC content | 1.02 | 0.34 | [0.33, 0.35] | 1.02 | 0.154 |
