## Supplementary methods for "Genomic insights into the karyotypic radiation of a narrow endemic holocentric plant *Carex helodes*"

**(a)**	**Experimental crosses, germination success rate estimation and cytogenetic study**

In spring 2019, experimental crosses involving the two populations were performed to generate a set of F₁ individuals. By late 2019, 50 achenes dry fruits from one cross were harvested and germinated under sterile conditions in a POL-EKO ST1 germinator, following an established protocol for *Carex helodes* [[1]](https://www.zotero.org/google-docs/?6RwV9z). Germinated seedlings were transplanted into pots and grown under greenhouse conditions. In spring 2021, once the F₁ plants reached reproductive maturity, self-fertilizations of F₁s were carried out to obtain an F₂ population. As in the previous generation, 50 achenes per self-cross (9 crosses in total) were tried to germinate under sterile conditions and seedlings were cultivated in the greenhouse until spring 2023. The germination success rate was calculated as the percentage of germinated seeds relative to the total number of seeds cultivated. The parents and two of the F₁ individuals from the crosses were karyotyped in spring 2021, when they started flowering. Anthers were fixed and stained [[2]](https://www.zotero.org/google-docs/?SwiHFn), and chromosomes and pairing were observed in metaphase I (MI) of meiosis using a Nikon Eclipse E400 microscope equipped with a digital camera Nikon DXM1200F. Five meiosis were counted per individual.

**(b) De novo repeat discovery and annotation**

Two methods were used to identify and annotate repetitive elements in the two genome assemblies. Repetitive regions were identified, classified and annotated using EarlGrey v4.4.5 [[3]](https://www.zotero.org/google-docs/?AR3oOQ). To further increase sensibility and minimize the amount of unclassified repetitive elements (REs) we used a RE library for plants including a library TE elements from TREP database [[4]](https://www.zotero.org/google-docs/?scCx64) and satellite library [[5]](https://www.zotero.org/google-docs/?FBvRwR) from PlantSat [[5]](https://www.zotero.org/google-docs/?Uo3N6U) completed with other REs identified in both assemblies through concatenated runs of EarlGrey with each assembly. To achieve this, Poales repetitive library was used to annotate and identify REs in AZN genome and the RE library obtained was then used in MON genome annotation, to obtain a final, common, *Carex helodes* RE library. Then, REs were filtered based on similarity and host genes were eliminated. This final library was used by RepeatMasker v4.1.5 [[6]](https://www.zotero.org/google-docs/?5nMIBb) to annotate repetitive elements in both genomes.

We complemented the previously described approach with more annotation tools specialized in different RE types with higher specificity. Transposable elements were further analyzed with DANTE [[7]](https://www.zotero.org/google-docs/?Vue8qA), available from the RepeatExplorer2 Galaxy portal (https://repeatexplorer-elixir.cerit-sc.cz/), exploiting the REXdb database (Viridiplantae [[8]](https://www.zotero.org/google-docs/?ZJBYW6)). An even more specific variation of the previous tool focused on LTR retrotransposons was utilized: DANTE_LTR [[7]](https://www.zotero.org/google-docs/?JPxcl2). From this portal, TAREAN annotation tool was also used, which focuses on tandem repeats which are commonly associated with holocentromeres [[9]](https://www.zotero.org/google-docs/?g6IAZi).

**(c) CRs validation and inversions heterozygosity**

Validation of chromosomal rearrangements was performed through: reciprocal Hi-C cross-mapping. Raw Hi-C reads from the AZN genome were mapped onto the MON reference genome and vice versa, using bwa mem v0.7.15 (Li & Durbin, 2009). Alignments were name-sorted, filtered for primary paired alignments, and converted into Juicer-compatible short format using custom scripts. Finally, cross-species Hi-C contact maps were generated using Juicer Tools v1.22.01 (Durand et al., 2016). These reciprocal contact maps enable the direct visualization of structural discordances using the JuiceBox 2.15 [72] software. To further confirm CRs, long-read split-read support was estimated for breakpoints predicted by GENESPACE following the pbsv pipeline [[10]](https://www.zotero.org/google-docs/?IYyzzQ), which was designed for PacBio data. Firstly, AZN HIFi reads were aligned against MON genome and vice versa using pbmm2 v26.1.99 [[11]](https://www.zotero.org/google-docs/?ja0eMI). Then, signatures of structural variation were detected and structural variants identified using the functions *‘discover’* and ‘*call*’ respectively from the pbsv software v2.11.0 [[10]](https://www.zotero.org/google-docs/?ygOQA0). Finally, breakends (BNDs) were searched within the breakpoints and their proximities (5000 bp to each side) and their support assessed through the number and proportion of split reads supporting the structural variant. In addition, inversion heterozygosity was assessed through the proportion of split-read support. Heterozygous inversions were confirmed through examining assembly Hi-C contact maps.
